# Gingipain-containing products from *Porphyromonas gingivalis* promote epithelial CCL20 signaling and γδ T-cell accumulation in COPD-like airways

**DOI:** 10.64898/2026.06.29.734663

**Authors:** Keisuke Kawano, Noriki Takahashi, Tomoki Kishimoto, Toru Kariu, Yukio Fujiwara, Mai Uemura, Kota Nakajima, Nodoka Kinjo, Keiko Ueno-Shuto, Ryunosuke Nakashima, Megumi Hayashi, Mary Ann Suico, Tsuyoshi Shuto

## Abstract

Chronic obstructive pulmonary disease (COPD) is a progressive inflammatory airway disease in which impaired mucosal barrier function may increase susceptibility to aspirated oral microbial products. Periodontal disease has been associated with COPD development and exacerbation, but the epithelial mechanisms linking periodontal pathogens to pulmonary immune remodeling remain unclear. Here, we investigated whether gingipain-containing *Porphyromonas gingivalis* culture supernatant (PCS) promotes γδ T-cell-associated inflammation in COPD-like airways. Repeated intratracheal administration of PCS to βENaC-transgenic mice induced airway-centered immune cell accumulation and increased γδ TCR-positive cell accumulation, together with elevated expression of the γδ T-cell-associated cytokines Ifng and Il17a. PCS also increased pulmonary *Ccl20* and *Ccr6* expression, whereas epithelial alarmin-related genes and M2 macrophage-associated responses were not induced in parallel. In ENaC-overexpressing human airway epithelial cells, PCS induced CCL20 and F2RL1, the gene encoding protease-activated receptor 2 (PAR-2), and reduced the N-terminal PAR-2 signal, consistent with proteolytic receptor cleavage. Direct PAR-2 activation reproduced CCL20 induction, whereas pharmacological PAR-2 inhibition suppressed PCS-induced CCL20 expression. In contrast, PAR-1 inhibition or LPS neutralization with polymyxin B did not suppress this response. These findings support a mucosal epithelial protease-sensing model in which gingipain-containing *P. gingivalis* products activate PAR-2-dependent CCL20 production in airway epithelial cells and are associated with CCR6-linked γδ T-cell accumulation in COPD-like airways.

**Contribution to the field statement:** Periodontal disease is often associated with chronic obstructive pulmonary disease, but how oral bacterial products affect lung inflammation remains unclear. This study shows that protease-rich products from the periodontal pathogen *Porphyromonas gingivalis* can be sensed by airway epithelial cells under COPD-like conditions. This response induces a chemokine signal that is linked to the accumulation of γδ T cells, a type of immune cell involved in inflammatory responses. Our findings suggest that the airway epithelium acts as an immune sensor, converting signals from aspirated oral bacterial protease into lung inflammation. This work provides a mechanistic framework for understanding how oral dysbiosis may contribute to immune remodeling in COPD.

## Introduction

Chronic obstructive pulmonary disease (COPD) is a progressive inflammatory airway disease characterized by mucus hypersecretion, small-airway remodeling, emphysematous destruction, and persistent airflow limitation. Long-term cigarette smoking is the major etiologic factor, and COPD remains a leading cause of morbidity and mortality worldwide (1–3). Chronic exposure to noxious stimuli injures airway epithelial cells, disrupts epithelial barrier function, and increases cytokine and chemokine production that recruits and activates inflammatory cells (4). Activated leukocytes, in turn, release reactive oxygen species, matrix metalloproteinases, and neutrophil elastase, thereby promoting protease-antiprotease imbalance, alveolar destruction, and disease progression (4, 5).

COPD frequently coexists with systemic comorbidities, particularly in older patients (6, 7). Among these conditions, periodontal disease has emerged as a potential independent risk factor for COPD development and exacerbation (8–10). Periodontal disease is a chronic infectious and inflammatory disease driven by dysbiotic bacterial colonization of periodontal tissues. *Porphyromonas gingivalis* (*P. gingivalis*), *Tannerella forsythia*, and *Treponema denticola*, collectively referred to as the red complex, are key periodontal pathogens (11). Oral bacteria and their products can enter the lower respiratory tract through aspiration, and *P. gingivalis* has been detected in bronchial samples from patients with COPD, particularly during acute exacerbations (12, 13). Moreover, the abundance of oral *P. gingivalis* negatively correlates with pulmonary function (12), supporting the possibility that recurrent aspiration of *P. gingivalis*-derived factors contributes to COPD pathogenesis (14).

Animal models of periodontitis-associated COPD have shown that *P. gingivalis* exposure augments pulmonary inflammation and worsens respiratory function compared with COPD alone (15, 16). In these models, γδ T cells accumulate in the lung and are accompanied by increased expression of inflammatory cytokines, including interferon-γ and interleukin-17 (15). M2 macrophage polarization has also been reported, suggesting that γδ T-cell-associated cytokine networks may amplify inflammatory and tissue-remodeling responses (15). However, the upstream mechanism by which *P. gingivalis* promotes γδ T-cell infiltration in COPD-like lungs remains incompletely defined.

*P. gingivalis* produces several virulence factors, including lipopolysaccharide, outer membrane vesicles, and cysteine proteases known as gingipains (17). Gingipains have been detected in lung tissues following *P. gingivalis* infection and have been implicated in *P. gingivalis*-induced pulmonary inflammation *in vivo* (18, 19). These proteases also contribute to systemic inflammatory diseases, including Alzheimer’s disease and nonalcoholic steatohepatitis (20, 21). Because airway epithelial cells form the first cellular barrier to aspirated microbial products and regulate leukocyte trafficking through chemokine production, we hypothesized that *P. gingivalis*-derived gingipains contribute to γδ T-cell accumulation through epithelial protease-sensing pathways.

In addition to conventional elastase-induced COPD models, our laboratory established an epithelial sodium channel β subunit (βENaC)-transgenic mouse model in which βENaC is overexpressed in airway epithelial cells (22). Elastase-induced COPD models develop acute protease-driven inflammation followed by emphysematous destruction of alveolar walls. In contrast, βENaC-transgenic mice develop airway surface liquid depletion, impaired mucociliary clearance, mucus accumulation, chronic immune cell infiltration, and progressive emphysema (23–26). These features recapitulate clinically relevant aspects of COPD, including small-airway mucus obstruction and increased βENaC expression in moderate-to-severe disease (27, 28).

In the present study, we used PCS, defined as a gingipain-containing extracellular protein fraction prepared from *P. gingivalis* culture supernatant, to determine whether periodontal bacterial protease activity is associated with γδ T-cell accumulation in COPD-like airways. Because PCS is a crude bacterial-derived preparation, we interpreted the in vivo responses as effects of gingipain-containing *P. gingivalis* products rather than as effects of purified gingipains alone.

## Materials and methods

### Animals

Wild-type C57BL/6J mice and C57BL/6J-βENaC-transgenic mice were used in this study. The βENaC-transgenic mouse model was established as described previously (22). B6.Cg-Tg(Scgb1a1-Scnn1b)6608Bouc/J mice were originally obtained from The Jackson Laboratory (Bar Harbor, ME, USA) and backcrossed with wild-type C57BL/6J mice. C57BL/6J-βENaC-transgenic mice generated from cryopreserved embryos or maintained by in-house breeding were used for the experiments. Mice were housed under specific pathogen-free conditions with a 12-h light/dark cycle at 20°C and were provided food and water ad libitum. Male mice were used at 9 weeks of age unless otherwise indicated.

All animal procedures were approved by the Animal Welfare Committee of Kumamoto University (#A2024-052) and conducted in accordance with institutional and national guidelines and ARRIVE recommendations.

### Preparation of *P. gingivalis* culture supernatant

*P. gingivalis* culture supernatant (PCS) was prepared as described previously (29). This preparation contains extracellular vesicle-associated bacterial components, including outer membrane vesicles, and closely corresponds to the preparation referred to as *P. gingivalis* outer membrane vesicles (Pg-OMVs). Briefly, *P. gingivalis* strain ATCC 33277 was cultured under anaerobic conditions in brain heart infusion medium supplemented with hemin and menadione. Bacterial cultures were centrifuged, and the resulting culture supernatants were collected. The protein fraction was concentrated by ammonium sulfate precipitation and dialyzed against phosphate buffer with saline to obtain a *P. gingivalis* extracellular protein/protease fraction containing gingipains; this preparation was defined as PCS. Protein concentration was determined before use. Because PCS was prepared from culture supernatant by ammonium sulfate precipitation and dialysis, it represents a concentrated extracellular protein/protease fraction rather than unfractionated bacterial culture supernatant. Therefore, PCS may contain gingipains together with other soluble or vesicle-associated *P. gingivalis*-derived components.

### Gingipain activity assay

Arginine-specific gingipain (Rgp) and lysine-specific gingipain (Kgp) activities in PCS were measured using fluorogenic substrate assays. PCS was diluted to the indicated concentrations in gingipain assay buffer consisting of 100 mM Tris-HCl (pH 7.5), 50 mM NaCl, and 5 mM CaCl2. L-cysteine was added to the buffer at a final concentration of 10 mM. Samples were incubated in a 96-well microplate at 37°C for 10 min.

For Rgp activity, Boc-Phe-Ser-Arg-MCA (Peptide Institute, Osaka, Japan, 3107-v) was used as the fluorogenic substrate. For Kgp activity, Boc-Val-Leu-Lys-MCA (Peptide Institute, 3104-v) was used. Each substrate was dissolved in DMSO at 5 mM and added to the reaction mixture at a final concentration of 50 μM. Fluorescence was measured immediately after substrate addition and again after incubation at 37°C for 30 min using a SpectraMax iD3 microplate reader (Molecular Devices, San Jose, CA, USA). The excitation and emission wavelengths were 365 and 460 nm, respectively. Gingipain activity was calculated as the difference in fluorescence intensity between 0 and 30 min.

### Intratracheal PCS administration

To model repeated aspiration of *P. gingivalis*-derived products, wild-type and βENaC-transgenic mice were administered PCS intratracheally. Immediately before administration, dithiothreitol (DTT), a reducing agent, was added to PCS at a final concentration of 1 mM. Mice were anesthetized with 3.5% isoflurane, and PCS was administered using a 1-mL Hamilton syringe connected to an oral gavage needle. PCS was administered daily at 400 μg/kg for 29 consecutive days. Control mice received vehicle containing the same final concentration of DTT. On day 30, pulmonary function was assessed, and lung tissues were collected for histological analysis, immunohistochemistry, and RNA isolation.

### Pulmonary function analysis

Pulmonary function was assessed using the flexiVent system (SCIREQ Inc., Montreal, QC, Canada). Mice were anesthetized by intraperitoneal injection of a mixed anesthetic solution containing medetomidine hydrochloride, midazolam, and butorphanol tartrate at doses of 0.75, 4, and 5 mg/kg, respectively. The anesthetic mixture was administered at 15 mL/kg.

After anesthesia, the trachea was exposed, incised, and cannulated with an 18-gauge needle. The cannula was secured by ligation, and mice were connected to the flexiVent system. Pulmonary mechanics, including resistance, elastance, and compliance, were measured. Forced expiratory volume in 0.1 s (FEV0.1) and forced vital capacity (FVC) were also measured, and FEV0.1/FVC was calculated as an index of airflow limitation.

### Tissue preparation

After pulmonary function analysis, the airway and whole lungs were excised and washed with PBS. The left lung was immersed in 10% neutral buffered formalin and fixed at room temperature for 12–24 h. After fixation, the left lung was divided into three parts and washed three times with PBS. Tissues were dehydrated with ethanol and subsequently processed for paraffin embedding by K.I. Stainer Co., Ltd (Kumamoto, Japan). Paraffin-embedded lung tissues were sectioned for histological and immunohistochemical analyses.

### Alcian blue/periodic acid-Schiff staining

Paraffin-embedded lung sections were deparaffinized with xylene and rehydrated through graded ethanol. For Alcian blue staining, sections were incubated in 3% acetic acid for 5 min and then stained with Alcian blue solution (pH 2.5; Nacalai Tesque, Kyoto, Japan) for 2 h at room temperature. After washing with 3% acetic acid, periodic acid–Schiff staining was performed using a Periodic Acid-Schiff staining kit according to the manufacturer’s protocol. Sections were incubated with periodic acid solution and Schiff’s reagent for 10–15 min each at room temperature. Nuclei were counterstained with hematoxylin. Sections were dehydrated through graded ethanol, cleared with xylene, and mounted with Canada balsam. Images were acquired using a BZ-X710 all-in-one fluorescence microscope (Keyence, Osaka, Japan).

### Mean linear intercept analysis

Emphysematous changes were evaluated by measuring the mean linear intercept (MLI), an index of alveolar airspace enlargement. Ten representative fields were selected from each lung section, including three upper, four middle, and three lower lung fields. A 300-μm square grid was superimposed on each field. The number of intersections between the grid lines and alveolar walls was counted, and MLI was calculated by dividing the total length of the grid lines by the number of intersections.

### Immunohistochemistry

Paraffin-embedded lung sections were deparaffinized with xylene and rehydrated through graded ethanol. Antigen retrieval was performed using Target Retrieval Solution. Endogenous peroxidase activity was quenched by incubation with 3% H_2_O_2_ in PBS or H_2_O_2_ in methanol for 15 min. After washing with PBS, sections were incubated overnight at 4°C with primary antibodies diluted in Antibody Diluent (DAKO, Glostrup, Denmark).

The following primary antibodies were used: rabbit anti-CD3 (Nichirei Biosciences, Tokyo, Japan, 413591), rabbit anti-CD4 (Cell Signaling Technology, Danvers, MA, USA, 25229), rabbit anti-CD8 (Cell Signaling Technology, Danvers, MA, USA, 98941), rabbit anti-CD206 (Abcam, Cambridge, UK, ab64693), rabbit anti-Iba1 (FUJIFILM Wako Pure Chemical Corporation, Osaka, Japan, 019-19741), rat anti-CD138 (BD Pharmingen^TM^, San Diego, CA, USA, 553712), and rabbit anti-Ly6G (Abcam, Cambridge, UK, ab238132). For γδ TCR staining, anti-γδ TCR antibody (LSBio, Shirley, MA, USA, LS-B5684) was used.

After three washes with PBS, sections were incubated with Histofine Simple Stain Mouse MAX-PO(R) (Nichirei Biosciences, Tokyo, Japan) for 1 h at room temperature. Signals were developed using DAB substrate (Nichirei Biosciences, Tokyo, Japan, 425011), and sections were counterstained with hematoxylin. Images were acquired using a NanoZoomer slide scanner (Hamamatsu Photonics, Hamamatsu, Japan).

### Quantification of immunohistochemical staining

Immunohistochemical staining was quantified using HALO image analysis software (Indica Labs, Albuquerque, NM, USA). Whole-lung section images acquired with the NanoZoomer scanner were imported into HALO, and image resolution was adjusted. Representative color tones for hematoxylin-positive nuclei and DAB-positive cells were registered for each slide, and threshold settings were optimized using the real-time tuning function. After confirming accurate recognition of DAB-positive cells, the entire slide was analyzed under the registered settings. The percentage of DAB-positive cells among total cells was calculated for each marker.

### Cell culture

Human bronchial epithelial 16HBE14o- cells and β/γENaC-overexpressing 16HBE14o- cells were used as normal and COPD model airway epithelial cells, respectively. 16HBE14o- cells were kindly provided by Dr. Dieter Gruenert, University of California, San Francisco. β/γENaC-overexpressing 16HBE14o- cells were kindly provided by Dr. Michael Stutts at the University of North Carolina and were used as previously described (30). Cells were cultured in MEM (FUJIFILM Wako Pure Chemical Corporation, Osaka, Japan) supplemented with 10% FBS (HyClone, Logan, UT, USA) and 1% penicillin/streptomycin (FUJIFILM Wako Pure Chemical Corporation) at 37°C in a humidified atmosphere containing 5% CO_2_. Culture dishes were coated with fibronectin coating solution before cell seeding. The coating solution consisted of 0.1 mg/mL MgCl_2_, 0.1 mg/mL CaCl_2_, 0.1 mg/mL BSA, 30 μg/mL collagen, and 10 μg/mL human plasma fibronectin in PBS containing MgCl_2_ and CaCl_2_. Coated dishes were incubated at 37°C for at least 30 min before use.

### Cell stimulation and inhibitor treatment

For PCS stimulation, 16HBE14o- cells and β/γENaC-overexpressing 16HBE14o- cells were treated with PCS at 300 ng/mL for 2 h for mRNA analysis or for 6 h for immunoblotting. PCS was diluted in culture medium immediately before use, and DTT was added at a final concentration of 10 μM. For direct PAR-2 activation, cells were treated with the PAR-2-activating peptide 2-furoyl-LIGRLO-NH_2_ (2-FLI; 300 nM) for 2 h. For inhibitor experiments, β/γENaC-overexpressing 16HBE14o- cells were preincubated with AZ3451, vorapaxar, or the LPS-neutralizing agent polymyxin B for 1 h before PCS stimulation. For experiments, AZ3451 and vorapaxar were diluted in culture medium to a final concentration of 10 μM, and polymyxin B was diluted to a final concentration of 10 μg/mL to neutralize LPS activity.After pretreatment, cells were stimulated with PCS at 300 ng/mL for 2 h.

### RNA isolation

Total RNA was isolated from mouse lung tissues and cultured cells using RNAiso Plus (Takara Bio Inc., Kusatsu, Shiga, Japan) according to the manufacturer’s protocol. For lung tissue RNA extraction, the left lung was excised, washed with PBS, immersed in RNAlater solution, and mixed overnight at 4°C. RNAlater solution was then removed, RNAiso Plus (Takara Bio Inc.) was added, and the tissue was homogenized using an overhead stirrer set (WHEATON, Millville, NJ, USA).

For cultured cells, cells were grown to near-confluence, washed twice with PBS, and lysed with RNAiso Plus (Takara Bio Inc.). All procedures were performed under RNase-free conditions. Chloroform was added at one-fifth the volume of RNAiso Plus (Takara Bio Inc.), and samples were vortexed for 10 s and incubated on ice for 3 min. After centrifugation at 12,000 rpm for 15 min at 4°C, the aqueous phase was transferred to a new tube. A second chloroform extraction was performed, followed by isopropanol precipitation. RNA pellets were washed twice with 75% ethanol, air-dried, and dissolved in 20 μL of DEPC-treated water.

For RNA extracted from mouse lung tissues, DNase treatment was performed according to the manufacturer’s protocol, followed by phenol/chloroform extraction and ethanol precipitation. RNA concentration and purity were measured using an Epoch spectrophotometer (BioTek Instruments, Winooski, VT, USA). RNA samples with an OD_260_/OD_280_ ratio greater than 1.8 were used for subsequent analyses.

### Quantitative real-time RT-PCR

cDNA was synthesized using PrimeScript RT Master Mix (Perfect Real Time; Takara Bio Inc.) according to the manufacturer’s protocol. Total RNA was used at 0.0625 μg per gene. Reverse transcription was performed using a LifeECO Thermal Cycler (Bioer Technology, Hangzhou, China) under the following conditions: 37°C for 30 min, 85°C for 10 s, and 4°C indefinitely. Quantitative real-time PCR was performed using TB Green Premix Ex Taq II (Tli RNaseH Plus; Takara Bio Inc.) and gene-specific primers on a CFX Connect Real-Time PCR Detection System (Bio-Rad, Hercules, CA, USA). The PCR cycling conditions were as follows: 95°C for 3 min, followed by 40 cycles of 95°C for 10 s and 65°C for 1 min. Gene expression was normalized to 18S rRNA or Gapdh as the internal control, as appropriate. Relative mRNA expression was calculated using the ΔΔCt method. Primer sequences are listed in Table 1.

**Table 1.**
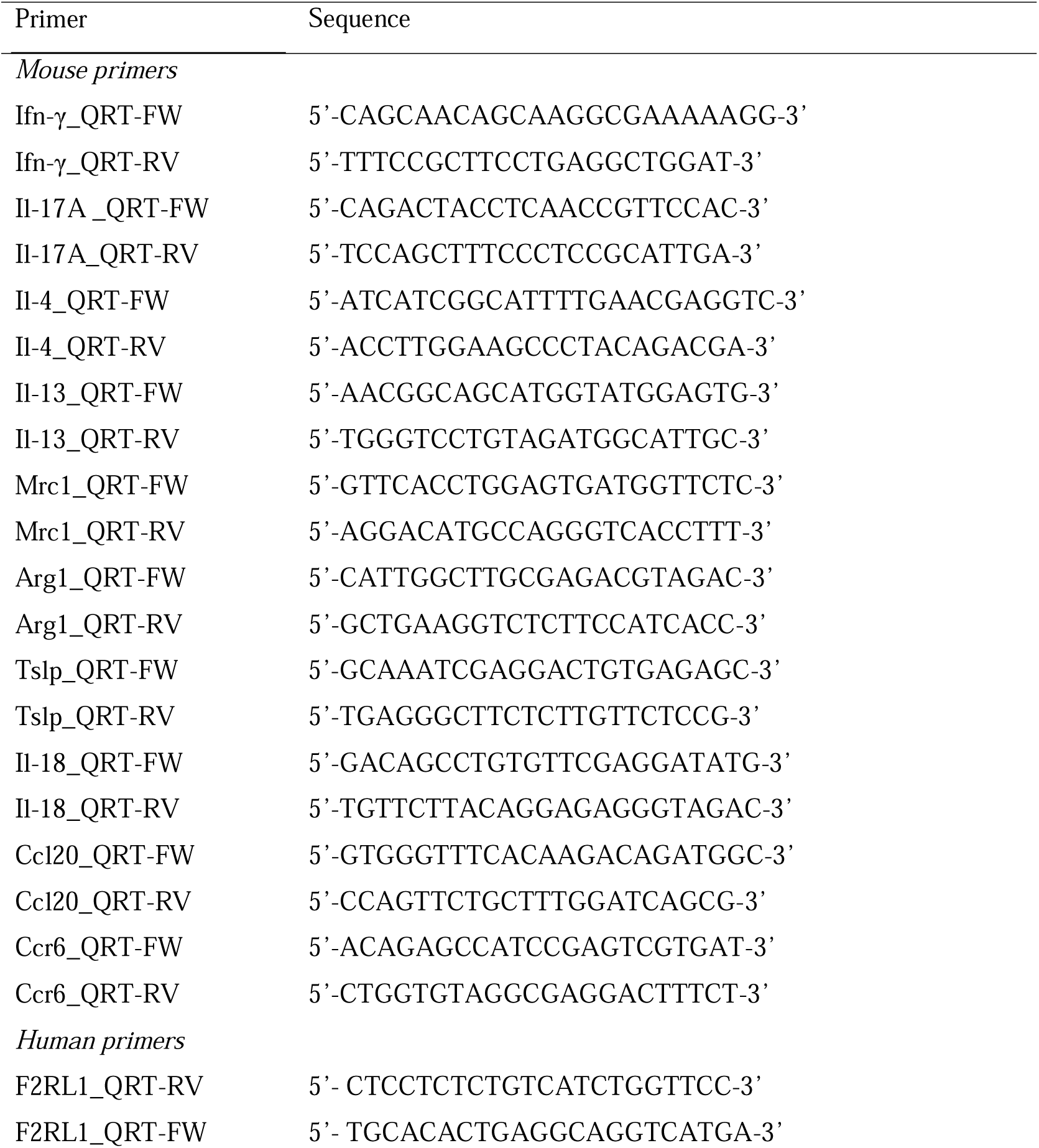
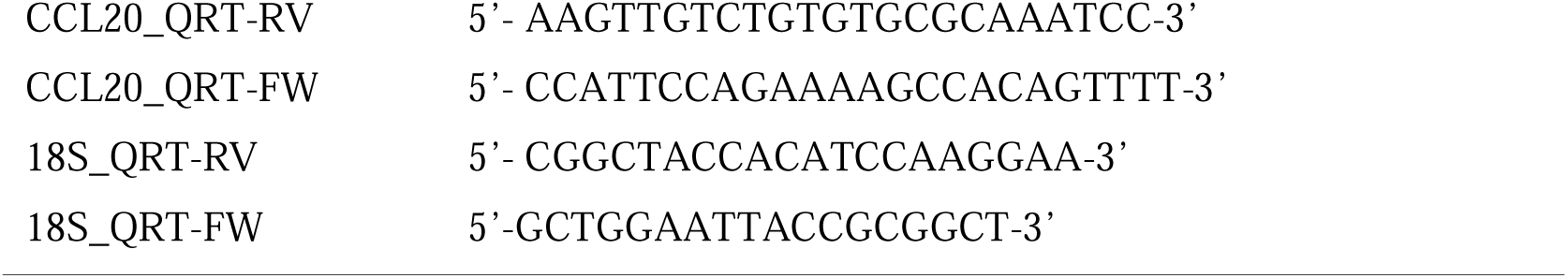
Primer sequences used for qRT-PCR.

### Immunoblotting

Cultured cells were grown to near-confluence, washed twice with PBS, and lysed in radioimmunoprecipitation assay buffer containing 50 mM Tris-HCl (pH 7.5), 150 mM NaCl, 1 mg/mL sodium deoxycholate, 1 mM Na_3_VO_4_, and 1% protease inhibitor cocktail. Cells were collected using a cell scraper and further lysed by pipetting. Lysates were incubated on ice for 30 min and centrifuged at 14,000 rpm for 30 min at 4°C. The supernatants were collected as protein extracts. Protein concentrations were determined using the bicinchoninic acid assay.

Equal amounts of protein were separated by 8% or 10% SDS-PAGE and transferred to PVDF membranes at 400 mA for 1 h. Membranes were blocked overnight at 4°C in 0.05% PBS-Tween containing 5% skim milk. After washing with 0.05% PBS-Tween, membranes were incubated with primary antibodies diluted in Can Get Signal Solution 1 for 2 h at room temperature. After washing, membranes were incubated with HRP-conjugated secondary antibodies diluted in Can Get Signal Solution 2 for 2 h at room temperature.

The following primary antibodies were used: rabbit anti-PAR-2 (Bioss Inc., Woburn, MA, USA, bs-1178R) and rat anti-HSC70 (Enzo Life Sciences, Farmingdale, NY, USA, ADI-SPA-815-D). The following secondary antibodies were used: anti-rabbit IgG-HRP and anti-rat IgG-HRP (Jackson ImmunoResearch, West Grove, PA, USA). Signals were detected using SuperSignal West Pico Chemiluminescent Substrate and visualized using an LAS-4000 mini imaging system (Fujifilm, Tokyo, Japan). Band intensities were quantified using Multi Gauge version 3.1 (Fujifilm) and normalized to HSC70.

### Statistical analysis

Data are presented as the mean ± SEM. Animal experiments included n = 5–7 mice per group, and cell-based experiments included n = 3–4 independent experiments, as indicated in the figure legends. Statistical analyses were performed using one-way ANOVA followed by the Tukey-Kramer multiple-comparison test. A value of *p* < 0.05 was considered statistically significant. All statistical analyses were performed using GraphPad Prism 10 (GraphPad Software, Boston, MA, USA).

## Results

### Intratracheal PCS administration induces airway-centered immune cell infiltration in COPD model mice

To model recurrent aspiration of *P. gingivalis*-derived products in periodontitis-associated COPD, wild-type and βENaC-transgenic mice received intratracheal PCS for 1 month at 400 µg/kg/day (Figure 1A). PCS was prepared from *P. gingivalis* strain ATCC 33277. Fluorogenic substrate assays confirmed concentration-dependent gingipain activity in PCS, with detectable arginine-specific gingipain and lysine-specific gingipain activities. Arginine-specific gingipain activity predominated over lysine-specific gingipain activity at each PCS concentration tested (Figure S1), consistent with previous reports that ATCC 33277 exhibits relatively high arginine-specific gingipain activity (31).

**Figure 1.**
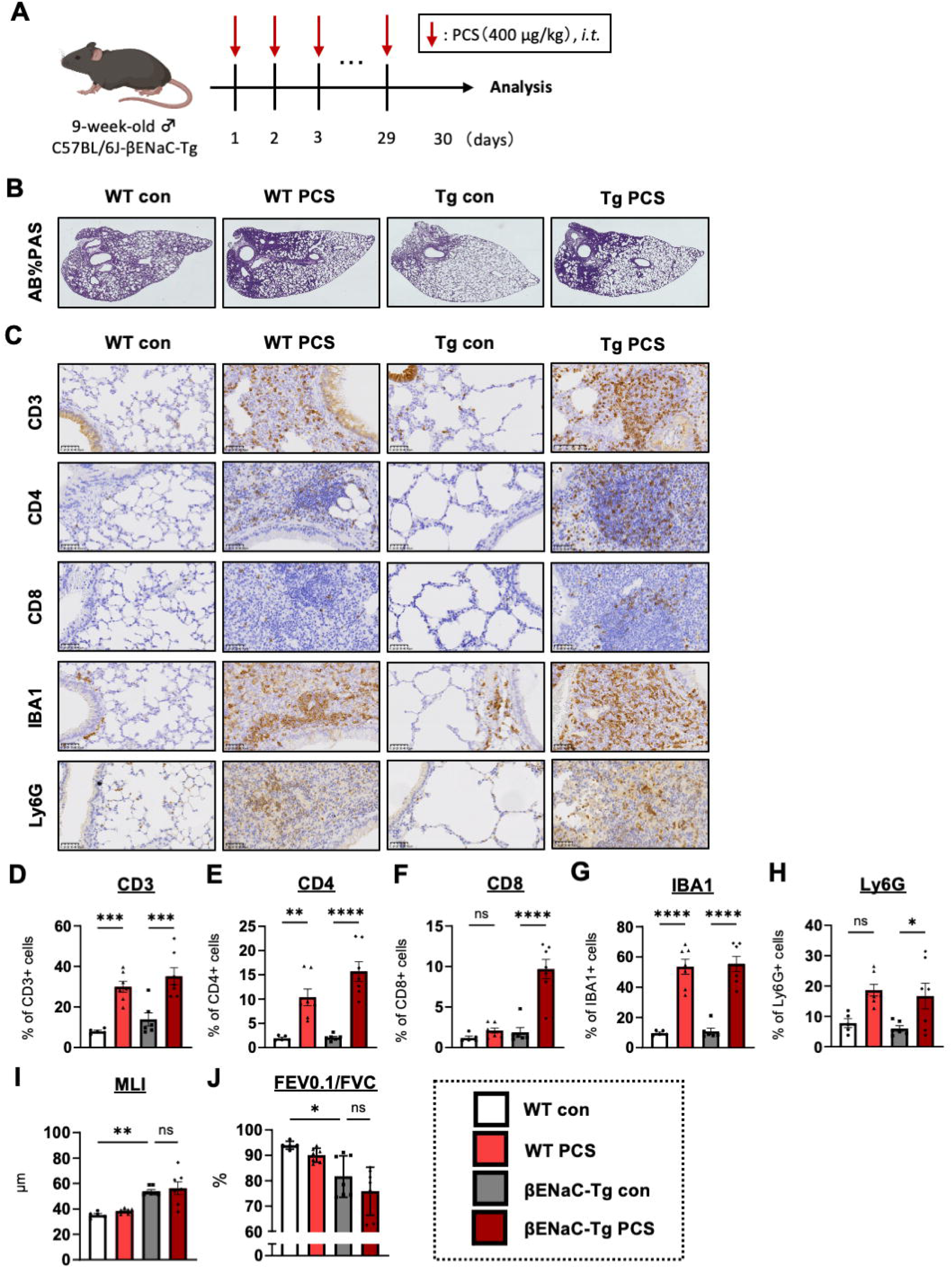
Intratracheal administration of PCS induces airway-centered immune cell infiltration in WT and. β**ENaC-Tg mice.** (A) Experimental scheme of repeated intratracheal administration of PCS to 9-week-old male C57BL/6J-βENaC-Tg mice. Mice received PCS at 400 µg/kg by intratracheal administration, and lung tissues and pulmonary function were analyzed on day 30. (B) Representative AB/PAS-stained lung sections from WT control, WT PCS-treated, βENaC-Tg control, and βENaC-Tg PCS-treated mice. (C) Representative immunohistochemical staining for CD3, CD4, CD8, IBA1, and Ly6G in lung sections from each group. (D–H) Quantification of CD3-positive cells (D), CD4-positive cells (E), CD8-positive cells (F), IBA1-positive cells (G), and Ly6G-positive cells (H), expressed as the percentage of marker-positive cells among total cells in the analyzed lung area. Quantification was performed using HALO image analysis software. (I) Quantitative morphometric analysis of emphysematous changes using the MLI. (J) Pulmonary function assessed as FEV0.1/FVC using the flexiVent system. Data are presented as the mean ± SEM; n = 5–7 mice per group. *P* values were assessed by one-way ANOVA followed by the Tukey-Kramer multiple-comparison test. \**p* < 0.05, \*\**p* < 0.01, \*\*\**p* < 0.001, and \*\*\*\**p* < 0.0001; ns, not significant.

PCS administration increased the accumulation of inflammatory cells around the airways in both wild-type and βENaC-transgenic mice. Immunohistochemical analysis showed increased numbers of CD3-, CD4-, and CD8-positive T cells, Ly6G-positive neutrophils, and IBA1-positive macrophages in PCS-treated lungs (Figures 1B–H). The increase in T-cell infiltration was more pronounced in βENaC-transgenic mice than in wild-type mice (Figures 1D–F). In contrast, PCS produced only modest changes in pulmonary mechanics and emphysematous morphology, as assessed by FEV0.1/FVC and alveolar septal morphometry (Figures 1I, J). These data indicate that PCS primarily induces airway-centered inflammatory cell infiltration under COPD-like airway conditions.

### PCS increases γδ T-cell accumulation and CCL20-CCR6 signaling without increasing M2 macrophage accumulation

We next examined whether PCS increases γδ T-cell accumulation in βENaC-transgenic lungs. Immunohistochemical staining for γδ TCR showed a marked increase in γδ T cells after 1 month of intratracheal PCS administration (Figures 2A, B). RT-qPCR analysis of whole-lung RNA showed increased expression of Ifng and Il17a, two major effector cytokines associated with γδ T-cell responses, in PCS-treated βENaC-transgenic mice (Figures 2C, D). Thus, PCS administration induced γδ T-cell accumulation together with an interferon-γ/interleukin-17A-associated inflammatory signature in COPD model lungs.

**Figure 2.**
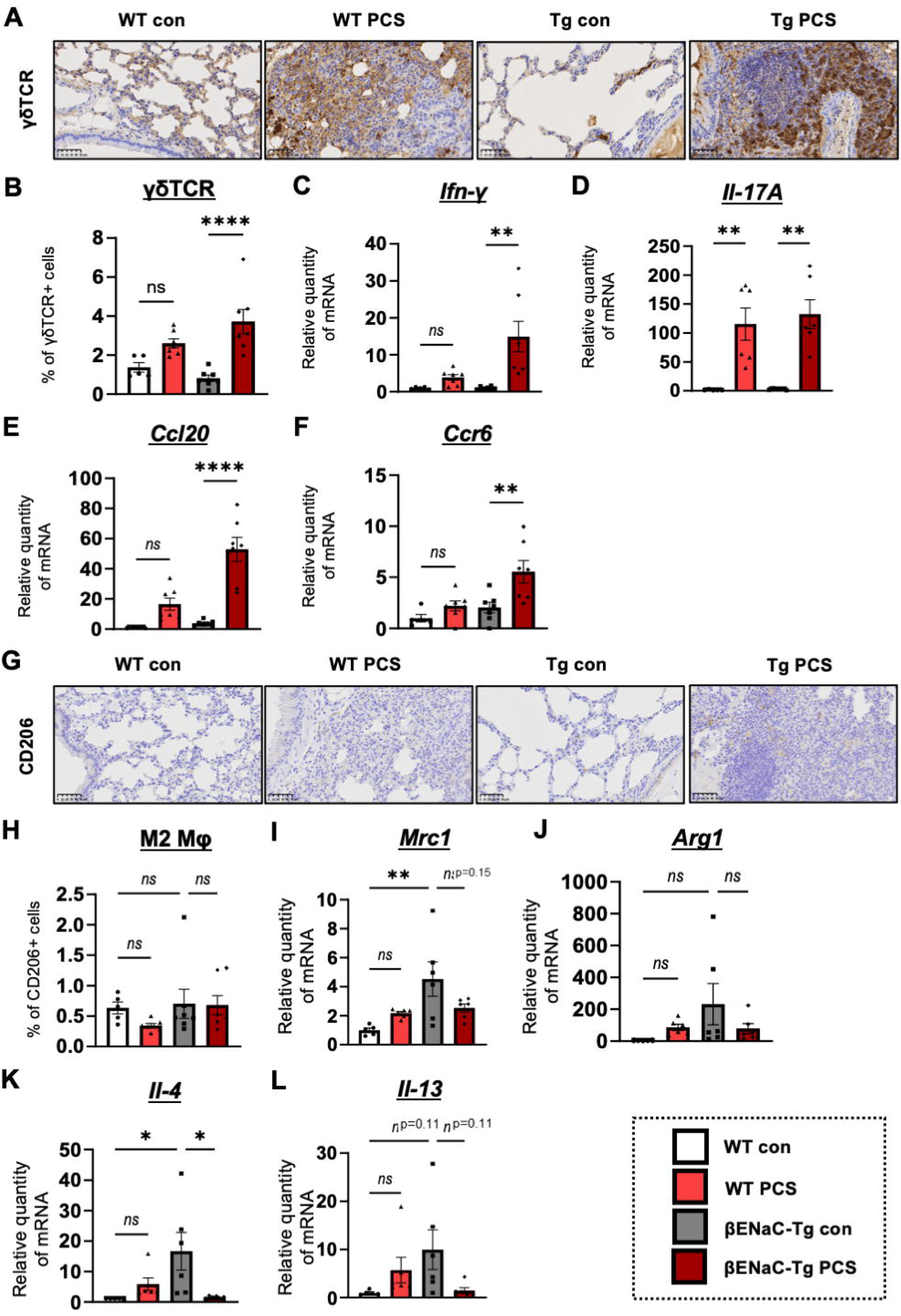
PCS increases γδ T-cell accumulation and CCL20-CCR6 signaling without increasing M2 macrophage accumulation in βENaC-Tg mouse lungs. (A) Representative immunohistochemical staining for γδ TCR in lung sections from WT control, WT PCS-treated, βENaC-Tg control, and βENaC-Tg PCS-treated mice. (B) Quantification of γδ TCR-positive cells, expressed as the percentage of marker-positive cells among total cells in the analyzed lung area. Quantification was performed using HALO image analysis software. (C–F) Relative mRNA expression of Ifn-γ (C), Il-17A (D), Ccl20 (E), and Ccr6 (F) in lung tissues, as measured by RT-qPCR. (G) Representative immunohistochemical staining for CD206 in lung sections from each group. (H) Quantification of CD206-positive M2 macrophages, expressed as the percentage of marker-positive cells among total cells in the analyzed lung area. Quantification was performed using HALO image analysis software. (I–L) Relative mRNA expression of M2 macrophage-associated genes and cytokines, including *Mrc1* (I), *Arg1* (J), *Il-4* (K), and *Il-13* (L), in lung tissues, as measured by RT-qPCR. Data are presented as the mean ± SEM; n = 5–7 mice per group. *P* values were assessed by one-way ANOVA followed by the Tukey-Kramer multiple-comparison test. \**p* < 0.05, \*\**p* < 0.01, and \*\*\*\**p* < 0.0001; ns, not significant.

Because airway epithelial cells produce chemokines that regulate leukocyte trafficking, we assessed the expression of γδ T-cell-associated chemotactic factors in the same lung samples. *Ccl20* expression was strongly increased in PCS-treated βENaC-transgenic mice, and *Ccr6* expression was also elevated (Figures 2E, F). CCL20 is an epithelial-derived chemokine that recruits CCR6-expressing γδ T cells, including interleukin-17A-producing γδ T cells, to inflamed tissues (32, 33). These results suggest that activation of the CCL20-CCR6 chemokine axis may contribute to PCS-associated γδ T-cell accumulation in COPD-like lungs.

We also evaluated M2 macrophage accumulation by staining for CD206. In contrast to γδ T cells, CD206-positive macrophages were not increased by PCS administration in either wild-type or βENaC-transgenic mice (Figures 2G, H). Consistent with this observation, the expression of M2-polarizing cytokines, including Il4 and Il13, tended to decrease after PCS treatment (Figures 2I, J). These data indicate that PCS-induced inflammation in this model is characterized by γδ T-cell accumulation rather than by M2 macrophage accumulation.

We therefore examined CD138-positive cells in lung tissues as a marker of plasma cell-like immune cell accumulation. Immunohistochemical analysis showed that PCS administration significantly increased the percentage of CD138-positive cells in both wild-type and βENaC-Tg mice (Fig. S2A,B). This increase was observed in both genotypes, suggesting that PCS promotes CD138-positive cell accumulation independently of βENaC-driven COPD-like airway remodeling. Because CD138 can also be expressed by non-plasma cell populations in a tissue-dependent manner, these data were interpreted as evidence of increased CD138-positive cell accumulation rather than definitive mature plasma cell expansion. Together with the increase in γδ T cells and the absence of M2 macrophage accumulation, these findings indicate that PCS induces a distinct airway-centered immune response characterized by γδ T-cell infiltration and CD138-positive cell accumulation

### PCS enhances Th1-associated gene expression in COPD model lungs

The increase in CD4- and CD8-positive T cells in PCS-treated βENaC-transgenic lungs suggested that PCS may alter T-cell-associated inflammatory programs. We therefore analyzed the expression of cytokines and transcription factors involved in Th1, Th2, Th17, and regulatory T-cell differentiation in whole-lung RNA samples (Figure S3). PCS induced only modest transcriptional changes in wild-type mice. In contrast, several Th1-associated genes were more strongly upregulated in PCS-treated βENaC-transgenic mice than in the other groups. Th17-related genes showed limited changes across groups. Although γδ T-cell infiltration and *Il17a* expression were increased by PCS, whole-lung RNA analysis did not show broad induction of a canonical Th17 differentiation program (Figure S3). Th2-related genes also showed no consistent increase, in agreement with the lack of CD206-positive M2 macrophage accumulation and the absence of *Il4* or *Il13* induction. Regulatory T-cell-related genes showed partial changes in PCS-treated βENaC-transgenic mice, but the pattern was not uniform across markers. Together, these data show that PCS preferentially enhances Th1-associated inflammatory signaling in COPD-like lungs.

### PCS does not induce epithelial cytokines implicated in γδ T-cell activation in COPD model lungs

Because PCS increased γδ T-cell accumulation together with *Ccl20* and *Ccr6* expression, we next asked whether PCS also induced epithelial cytokines that can regulate γδ T-cell activation. Thymic stromal lymphopoietin is an epithelial cytokine induced by inflammatory stimuli and can enhance interleukin-17A production by γδ T cells indirectly through dendritic cell activation (34, 35). Interleukin-18, which can be released upon epithelial injury, promotes γδ T-cell expansion and activation and augments interferon-γ production (36, 37). Therefore, we examined whether repeated intratracheal administration of PCS altered *Tslp* or *Il18* expression in lung tissue.

RT-qPCR analysis showed that PCS did not significantly change the expression of either Tslp or Il18 in βENaC-transgenic mouse lungs (Figures S4A, B). These findings suggest that PCS-induced γδ T-cell accumulation is not accompanied by broad induction of epithelial cytokines involved in γδ T-cell activation. Rather, the response appears to be associated predominantly with activation of the CCL20-CCR6 chemotactic axis.

### PCS activates PAR-2 and induces CCL20 expression in COPD model airway epithelial cells

To identify an epithelial mechanism upstream of CCL20 induction, we used β/γENaC-overexpressing 16HBE14o- cells as a COPD model of airway epithelial cells (30). PCS treatment at 300 ng/mL for 2 h increased CCL20 mRNA expression, and this response was greater in β/γENaC-overexpressing cells than in parental 16HBE14o- cells (Figures 3A, B). PCS also increased F2RL1 mRNA expression, which encodes PAR-2 (Figure 3B).

**Figure 3.**
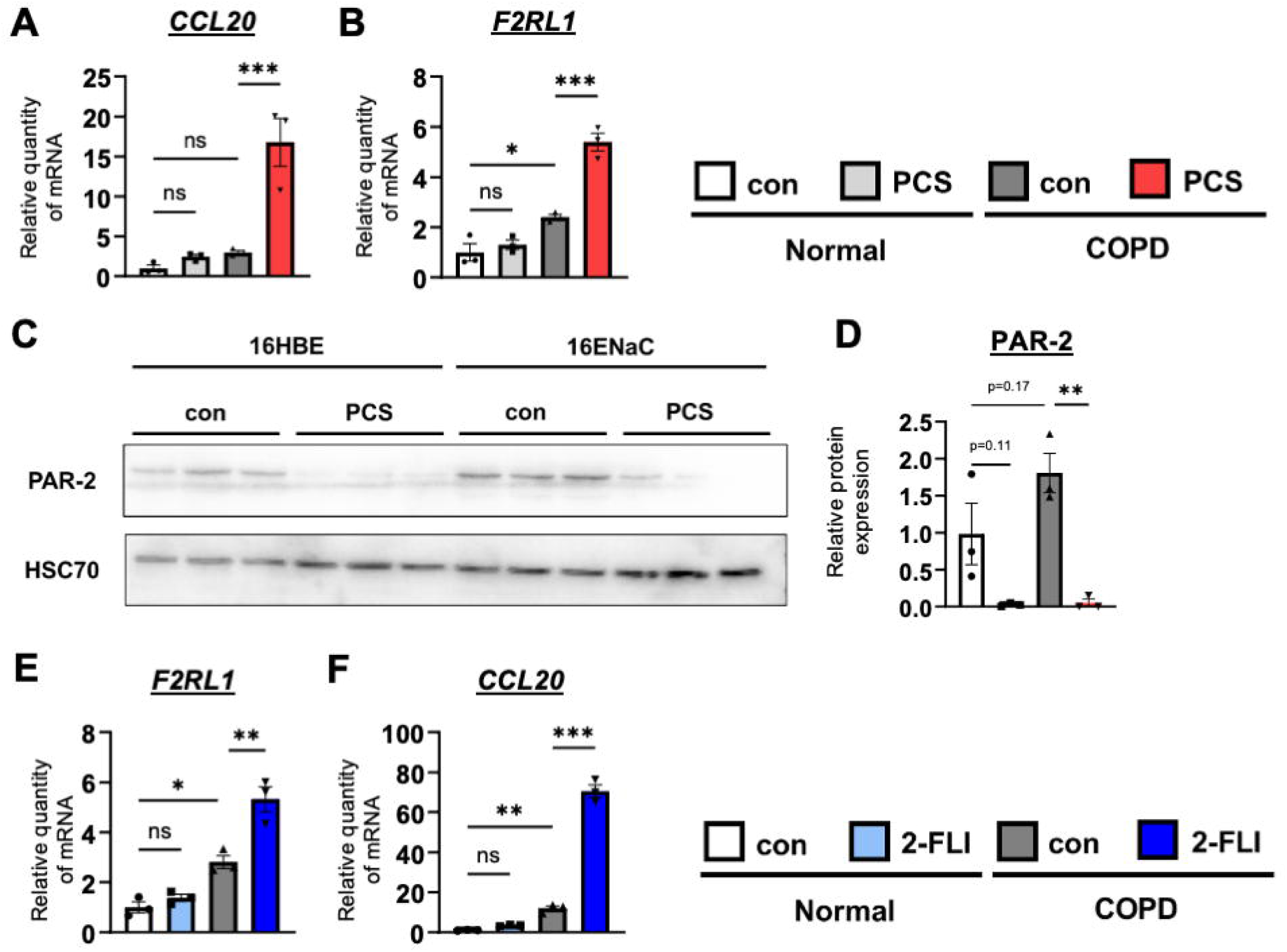
PCS and PAR-2 activation induce CCL20 expression in COPD model airway epithelial cells. (A, B) Normal human bronchial epithelial 16HBE14o- cells and COPD model β/γENaC-overexpressing 16HBE14o- cells were treated with PCS (300 ng/mL) for 2 h. Relative mRNA expression of CCL20 (A) and F2RL1 (B), which encodes PAR-2, was measured by quantitative real-time RT-PCR and normalized to 18S rRNA. (C, D) PAR-2 protein expression in 16HBE14o- and β/γENaC-overexpressing 16HBE14o- cells treated with PCS (300 ng/mL) for 6 h was analyzed by immunoblotting using an antibody recognizing the N-terminus of PAR-2. HSC70 was used as the loading control. Representative immunoblots are shown in (C), and densitometric quantification of PAR-2 protein expression normalized to HSC70 is shown in (D). (E, F) Cells were treated with the PAR-2-FLI (300 nM) for 2 h, and relative mRNA expression of F2RL1 (E) and CCL20 (F) was measured by quantitative real-time RT-PCR and normalized to 18S rRNA. Data are presented as the mean ± SEM; n = 3 independent experiments. *P* values were assessed by one-way ANOVA followed by the Tukey-Kramer multiple-comparison test. \**p* < 0.05, \*\**p* < 0.01, and \*\*\**p* < 0.001; ns, not significant.

Gingipains can activate PAR-2 by proteolytic cleavage of the receptor N-terminus (38, 39). Consistent with receptor cleavage, immunoblotting with an antibody recognizing the PAR-2 N-terminus showed loss of the PAR-2 band after PCS treatment at 300 ng/mL for 6 h (Figures 3C, D). Basal PAR-2 expression was also higher in β/γENaC-overexpressing cells than in parental cells, suggesting increased responsiveness of COPD model epithelial cells to protease stimulation.

We then tested whether direct PAR-2 activation reproduced the effect of PCS. Treatment with the PAR-2-activating peptide 2-furoyl-LIGRLO-NHS at 300 nM for 2 h increased both CCL20 and F2RL1 expression in β/γENaC-overexpressing 16HBE14o-cells (Figures 3E, F). These data show that PAR-2 activation is sufficient to induce the CCL20 response observed after PCS exposure in COPD model airway epithelial cells.

### PAR-2 inhibition suppresses PCS-induced CCL20 expression, whereas PAR-1 inhibition or LPS neutralization does not

To determine whether PAR-2 mediates PCS-induced CCL20 expression, β/γENaC-overexpressing 16HBE14o- cells were pretreated with the PAR-2 antagonist AZ3451 before PCS stimulation. PAR-2 inhibition markedly suppressed PCS-induced CCL20 expression (Figure 4A), demonstrating that PAR-2 activity is required for the epithelial CCL20 response to PCS.

**Figure 4.**
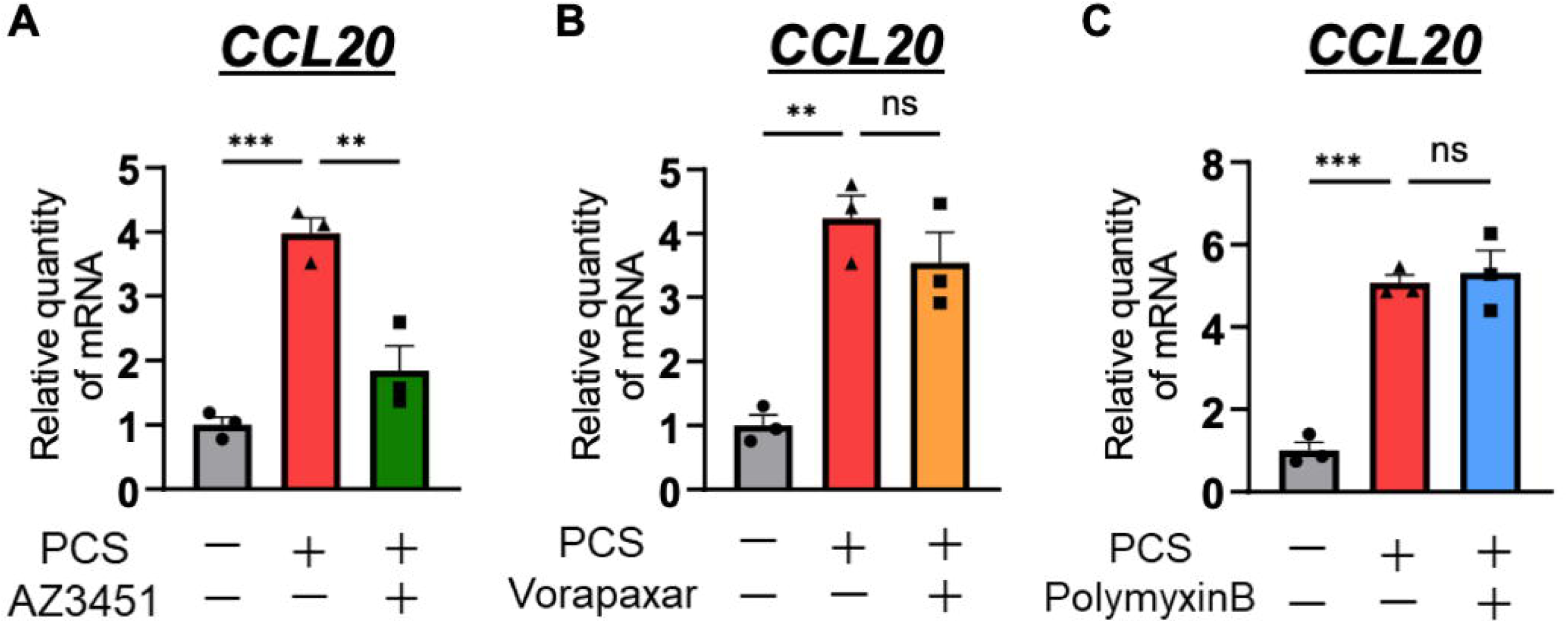
PAR-2 inhibition suppresses PCS-induced CCL20 expression, whereas PAR-1 inhibition or LPS neutralization does not. COPD model β/γENaC-overexpressing 16HBE14o- cells were preincubated with the PAR-2 antagonist AZ3451 (10 μM) (A), the PAR-1 antagonist vorapaxar (10 μM) (B), or the LPS-neutralizing agent polymyxin B (C) for 1 h, followed by stimulation with PCS (300 ng/mL) for 2 h. Relative mRNA expression of CCL20 was measured by quantitative real-time RT-PCR and normalized to 18S rRNA. Data are presented as the mean ± SEM; n = 3 independent experiments. *P* values were assessed by one-way ANOVA followed by the Tukey-Kramer multiple-comparison test. \*\**p* < 0.01 and \*\*\**p* < 0.001; ns, not significant.

Because gingipains can also activate PAR-1 and PCS may contain *P. gingivalis*-derived lipopolysaccharide, we tested whether PAR-1 activity or an LPS-dependent component contributes to PCS-induced CCL20 expression. Pretreatment with the PAR-1 antagonist vorapaxar or the LPS-neutralizing agent polymyxin B did not suppress PCS-induced CCL20 upregulation (Figures 4B, C). Thus, under these conditions, PCS-induced CCL20 expression was dependent on PAR-2 and was not detectably attenuated by PAR-1 inhibition or LPS neutralization.

## Discussion

This study identifies an airway epithelial signaling pathway by which gingipain-containing *P. gingivalis* culture supernatant promotes γδ T-cell-associated inflammation under COPD-like conditions. Repeated intratracheal administration of PCS to βENaC-transgenic mice increased γδ TCR-positive cell accumulation in the lung and elevated the expression of the γδ T-cell-associated cytokines Ifng and Il17a. This response was accompanied by increased expression of *Ccl20* and *Ccr6*, whereas the epithelial alarmin-related genes examined in this study did not show parallel induction. In contrast, PCS did not promote the accumulation of CD206-positive M2 macrophages or the expression of M2-polarizing cytokines. In the COPD model, airway epithelial cells, PCS increased CCL20 and F2RL1 expression and reduced the N-terminal PAR-2 signal, consistent with proteolytic cleavage of PAR-2. Pharmacological inhibition of PAR-2 suppressed PCS-induced CCL20 expression, whereas PAR-1 inhibition or LPS neutralization with polymyxin B did not. Moreover, direct stimulation with a PAR-2 agonist reproduced the induction of CCL20. Together, these findings support a model in which gingipain-containing *P. gingivalis* products activate PAR-2-dependent CCL20 expression in COPD-like airway epithelial cells. *In vivo*, PCS administration was associated with increased pulmonary *Ccl20* and *Ccr6* expression and accumulation of γδ TCR-positive cells, suggesting that this epithelial pathway may contribute to CCR6-linked γδ T-cell accumulation or enrichment in COPD-like airways.

The epithelial origin of this response is an important feature of the proposed mechanism. Airway epithelial cells are positioned at the interface between aspirated oral bacterial products and the lung immune environment, and they produce chemokines that shape leukocyte recruitment. CCL20 is particularly relevant because its receptor, CCR6, is expressed by interleukin-17-producing γδ T cells and has been implicated in γδ T-cell trafficking within inflamed tissues (40, 41). The increase in *Ccl20* and *Ccr6* expression observed in PCS-treated βENaC-transgenic lungs, together with the PAR-2-dependent induction of CCL20 in β/γENaC-overexpressing epithelial cells, links a bacterial protease signal to a chemokine pathway capable of explaining the γδ T-cell phenotype.

The absence of a clear *Tslp* or *Il18* response further suggests that PCS-induced γδ T-cell accumulation in this model is more closely associated with the CCL20-CCR6 axis than with a broad epithelial alarmin program.

These results extend previous work showing that periodontitis can aggravate COPD-like pathology through γδ T-cell activation and M2 macrophage polarization (15), but they also define an important difference between direct PCS exposure and whole *P. gingivalis*-associated disease models. In the present study, PCS increased γδ T-cell accumulation and induced an interferon-γ/interleukin-17A-associated inflammatory environment, but it did not recapitulate the marked M2 macrophage response or prominent deterioration in COPD-related pathological indices reported in periodontitis-COPD models. This difference likely reflects the fact that PCS exposure is not equivalent to chronic oral infection with viable *P. gingivalis*. Whole-bacterial infection may provide persistent microbial stimulation, periodontal tissue inflammation, repeated aspiration, outer membrane vesicles, LPS and other TLR ligands, short-chain fatty acids, and host immune priming, all of which could cooperate with gingipain-dependent epithelial signaling (32, 33, 42). Thus, gingipain-containing PCS appears to capture an epithelial chemokine module associated with γδ T-cell accumulation, whereas additional bacterial or host-derived signals may be required to drive the full pathological cascade leading to macrophage polarization and overt COPD exacerbation.

One possible explanation for the lack of M2 macrophage accumulation is that PCS shifted the lung immune environment away from a Th2/M2-polarizing state. In the present study, PCS increased the expression of Th1-associated genes in βENaC-transgenic lungs, whereas Th2-related genes were not consistently induced, and the expression of M2-polarizing cytokines, including *Il4* and *Il13*, tended to decrease. IL-4 and IL-13 are well-established drivers of alternative macrophage activation, whereas interferon-γ promotes classically activated inflammatory macrophage programs and can antagonize IL-4-dependent responses. Thus, the Th1-skewed inflammatory milieu induced by PCS may have limited IL-4/IL-13-dependent M2 macrophage differentiation, thereby explaining why PCS induced γδ T-cell infiltration without producing the robust M2 macrophage response observed in whole-*P. gingivalis* periodontitis-COPD models.

Differences in the protease composition of *P. gingivalis* strains may also contribute to distinct inflammatory outcomes. The PCS used in this study, derived from *P. gingivalis* ATCC 33277, exhibited predominant arginine-specific gingipain activity relative to lysine-specific gingipain activity. Because arginine-specific and lysine-specific gingipains differ in cleavage specificity and substrate preference, the balance between these protease activities may influence whether epithelial chemokine production, cytokine degradation, neutrophil recruitment, or macrophage polarization predominates (17, 43, 44). One possible explanation for the lack of M2 macrophage induction is that gingipain-rich PCS suppressed or degraded signals required for M2 polarization, such as IL-4 and IL-13, rather than promoting the cytokine environment observed in whole-infection models. This interpretation is consistent with the reduced or noninduced expression of M2-associated mediators in PCS-treated lungs, but it remains to be tested directly using purified arginine-specific and lysine-specific gingipains, gingipain-deficient bacterial products, and cytokine stability assays.

The COPD-like epithelial state may sensitize the lung to gingipain-PAR-2 signaling. βENaC-transgenic mice and β/γENaC-overexpressing airway epithelial cells model airway surface dehydration, impaired mucus clearance, and inflammatory epithelial dysfunction. In the present cell model, basal PAR-2 expression was higher in β/γENaC-overexpressing cells than in control epithelial cells, and PCS-induced CCL20 expression was more pronounced under this COPD-like condition. These findings suggest that ENaC-driven epithelial stress may lower the threshold for PAR-2-dependent inflammatory signaling. This interpretation is consistent with studies showing that protease-dependent PAR-2 signaling can amplify inflammatory responses in airway epithelial cells (45, 46). Because ENaC activity is regulated by proteolytic cleavage and can be activated by airway proteases, the ENaC-overexpressing epithelial state may be particularly susceptible to protease-rich microbial products (47–51).

The modest effect of PCS on lung function and emphysematous pathology should be interpreted in this mechanistic context. The present data indicate that γδ T-cell accumulation can be separated from overt M2 macrophage polarization and severe structural deterioration. Thus, γδ T-cell accumulation may represent an early or partial inflammatory response to gingipain-containing bacterial products rather than a sole driver of COPD progression. Additional stimuli, such as cigarette smoke exposure, elastase-driven injury, viable bacterial infection, or longer-term repeated aspiration, may be necessary to convert this epithelial chemokine response into persistent neutrophilic inflammation, macrophage remodeling, and measurable respiratory impairment. This distinction may also explain why different COPD models and different timing of protease exposure produce divergent outcomes.

The PCS-induced γδ T-cell response may also represent an early inflammatory step toward lymphoid organization rather than mature tertiary lymphoid tissue formation. Pulmonary tertiary lymphoid tissues, also referred to as inducible bronchus-associated lymphoid tissue, have been implicated in chronic lung inflammation and COPD, where persistent inflammatory stimulation promotes the local organization of B cells, T cells, dendritic cells, and lymphoid chemokines such as CXCL13, CCL19, and CCL21 (52,53). In addition, IL-17A has been linked to cigarette smoke-induced lymphoid neogenesis in COPD lungs (54,55). In this study, PCS increased γδ T-cell accumulation, *Il17a* expression, and CCL20-CCR6-associated signaling, but we did not assess B-cell follicles, CXCL13, CCL19, CCL21, high endothelial venules, or follicular dendritic cell networks. Therefore, the present findings do not demonstrate mature tertiary lymphoid tissue formation. Rather, they suggest that gingipain-dependent epithelial activation may initiate a chemokine-driven lymphocyte recruitment program that could precede lymphoid neogenesis under chronic COPD-like inflammatory conditions.

Several limitations should be considered. First, although PCS contained measurable gingipain activity and induced PAR-2-dependent CCL20 expression in airway epithelial cells, the present *in vivo* experiments do not establish gingipains as the sole active components of PCS. Because PCS was prepared as a concentrated extracellular protein/protease fraction from *P. gingivalis* culture supernatant, it may contain gingipains together with other bacterial products, including outer membrane vesicle-associated factors, LPS, and soluble virulence molecules. Although polymyxin B treatment did not suppress PCS-induced CCL20 expression, this result should be interpreted as evidence against a major LPS-dependent contribution under the present conditions rather than as definitive exclusion of all TLR4-dependent signaling. Future studies using purified Rgp and Kgp, gingipain-deficient *P. gingivalis* preparations, gingipain inhibitors, protease-inactivated PCS, more selective TLR4 pathway inhibitors such as TAK-242, or genetic approaches will be required to define the specific contribution of gingipain activity and TLR4-dependent signaling. Second, the causal role of the PAR-2-CCL20-CCR6 axis was demonstrated primarily in epithelial cell experiments and inferred *in vivo* from increased Ccl20 and Ccr6 expression, together with accumulation of γδ TCR-positive cells. *In vivo* blockade or genetic deletion of PAR-2, CCL20, or CCR6 will be necessary to determine whether this pathway is required for PCS-induced γδ T-cell accumulation in COPD-like lungs. Measurement of CCL20 protein levels in bronchoalveolar lavage fluid or lung homogenates would also strengthen the link between epithelial chemokine induction and immune cell accumulation. Third, γδ T-cell subsets and the cellular sources of CCL20 were not resolved in the present study. Flow cytometry, immunofluorescence colocalization, epithelial cell-specific analyses, or single-cell approaches will be needed to determine whether PCS preferentially enriches IL-17A-producing CCR6-positive γδ T cells and whether airway epithelial cells are the dominant source of CCL20 *in vivo*.

In summary, this study proposes that *P. gingivalis*-derived gingipain-containing products engage PAR-2 on COPD-like airway epithelial cells, thereby inducing CCL20 and promoting γδ T-cell-associated lung inflammation. The findings place the airway epithelium upstream of the γδ T-cell response and distinguish epithelial chemokine induction from the broader pathological program required for severe COPD exacerbation. These results support a model in which the gingipain-PAR-2-CCL20 axis serves as an inflammatory entry point that links exposure to periodontal bacterial proteases to immune remodeling in COPD-like airways, while full disease progression likely requires additional microbial and environmental signals.

## Data availability statement

The original contributions presented in the study are included in the article/Supplementary Material. The raw data supporting the conclusions of this article will be made available by the authors, without undue reservation.

## Ethics statement

All animal procedures were approved by the Animal Welfare Committee of Kumamoto University (#A2024-052) and conducted in accordance with institutional and national guidelines and ARRIVE recommendations.

## Abbreviations

CCL20: C-C motif chemokine ligand 20
COPD: chronic obstructive pulmonary disease
CCR6: C-C motif chemokine receptor 6
ENaC: epithelial sodium channel
Kgp: lysine-specific gingipain
PAR-1: protease-activated receptor 1
PAR-2: protease-activated receptor 2
PCS: *Porphyromonas gingivalis* culture supernatant
Rgp: arginine-specific gingipain
TLR4: Toll-like receptor 4.

## Acknowledgements

The authors thank the members of the Department of Molecular Medicine, Graduate School of Pharmaceutical Sciences, Kumamoto University, for technical assistance and helpful discussions. The authors would also like to express their sincere gratitude to Dr. Hirofumi Kai, Professor Emeritus of Kumamoto University, for his invaluable scientific guidance.

## Author contributions

K.K., N.T., and T.S. designed research. K.K., N.T. and K.U.-S. performed research and analyzed data. T.Ki., M.U., K.N., N.K., R.N., and M.H. contributed to the mouse experiments. K.K., N.T., and T.Ka. prepared gingipain-containing PCS. K.K., N.T., and Y.F. performed tissue staining and HALO calculation. K.K., N.T., M.A.S., and T.S. wrote the paper. T.S. supervised the project. All authors reviewed and edited the manuscript, contributed to the article, and approved the submitted version.

## Funding

This work was supported by Japan Society for the Promotion of Science (JSPS) KAKENHI (JP23K06150 to T.S.); the Program for Building Regional Innovation Ecosystems at Kumamoto University; the Health Life Science S-HIGO (Health life science: Interdisciplinary and Glocal Oriented) Professional Fellowship Program; the Program for Fostering Innovators to Lead a Better Co-being Society (JPMJSP2127; MEXT, Japan); and the Nagai Memorial Research Scholarship from the Pharmaceutical Society of Japan (N-217201 to N.T. and N-197203 to R.N.).

## Conflict of interest

The authors declare that the research was conducted in the absence of any commercial or financial relationships that could be construed as a potential conflict of interest.

## Supplementary material

The Supplementary Material for this article can be found online.

## Supplementary figure legends

**Figure S1. Concentration-dependent gingipain activity in PCS.**

Lysine-specific gingipain (Kgp) and arginine-specific gingipain (Rgp) activities in PCS were measured using fluorogenic substrate assays. PCS was tested at the indicated concentrations, and enzymatic activity is shown as fluorescence intensity. Data are presented as the mean ± SEM; n = 3 independent measurements.

**Figure S2. Intratracheal administration of PCS induces airway-centered CD138-positive cell accumulation in WT and βENaC-Tg mice.**

(A) Representative immunohistochemical staining for CD138 in lung sections from each group. (B) Quantification of CD138-positive cells expressed as the percentage of marker-positive cells among total cells in the analyzed lung area. Quantification was performed using HALO image analysis software. Data are presented as the mean ± SEM; n = 5–7 mice per group. P values were assessed by one-way ANOVA followed by the Tukey-Kramer multiple-comparison test. **p < 0.01, ****p < 0.0001.

**Figure S3. PCS preferentially enhances Th1-associated gene expression in βENaC-Tg mouse lungs.**

Relative mRNA expression of T-cell differentiation-related cytokines and transcription factors was measured by RT-qPCR in lung tissues from WT control, WT PCS-treated, βENaC-Tg control, and βENaC-Tg PCS-treated mice. Genes associated with Th1 responses (*Ifng*, *Il12a*, *Il12rb1*, and *Lta*), Th17 responses (*Il6*, *Tgfb*, and *Rorc*), Th2 responses (*Il4r*, *Il5*, *Il13*, and *Gata3*), and regulatory T-cell responses (*Il2*, *Il2ra*, and *Foxp3*) were analyzed. Data are presented as the mean ± SEM; n = 5–7 mice per group. *P* values were assessed by one-way ANOVA followed by the Tukey-Kramer multiple-comparison test. \**p* < 0.05, \*\**p* < 0.01, \*\*\**p* < 0.001, and \*\*\*\**p* < 0.0001; ns, not significant. Exact *p* values are shown where applicable.

**Figure S4. PCS does not significantly induce epithelial cytokines associated with γδ T-cell activation in βENaC-Tg mouse lungs.**

Relative mRNA expression of Tslp (A) and Il18 (B) was measured by RT-qPCR in lung tissues from WT control, WT PCS-treated, βENaC-Tg control, and βENaC-Tg PCS-treated mice. Data are presented as the mean ± SEM; n = 5–7 mice per group. *P* values were assessed by one-way ANOVA followed by the Tukey-Kramer multiple-comparison test. ns, not significant.

## Notes

### Competing Interest Statement

The authors have declared no competing interest.

