## Supplemental Figures for "Gingipain-containing products from *Porphyromonas gingivalis* promote epithelial CCL20 signaling and γδ T-cell accumulation in COPD-like airways"

### Figure.S1

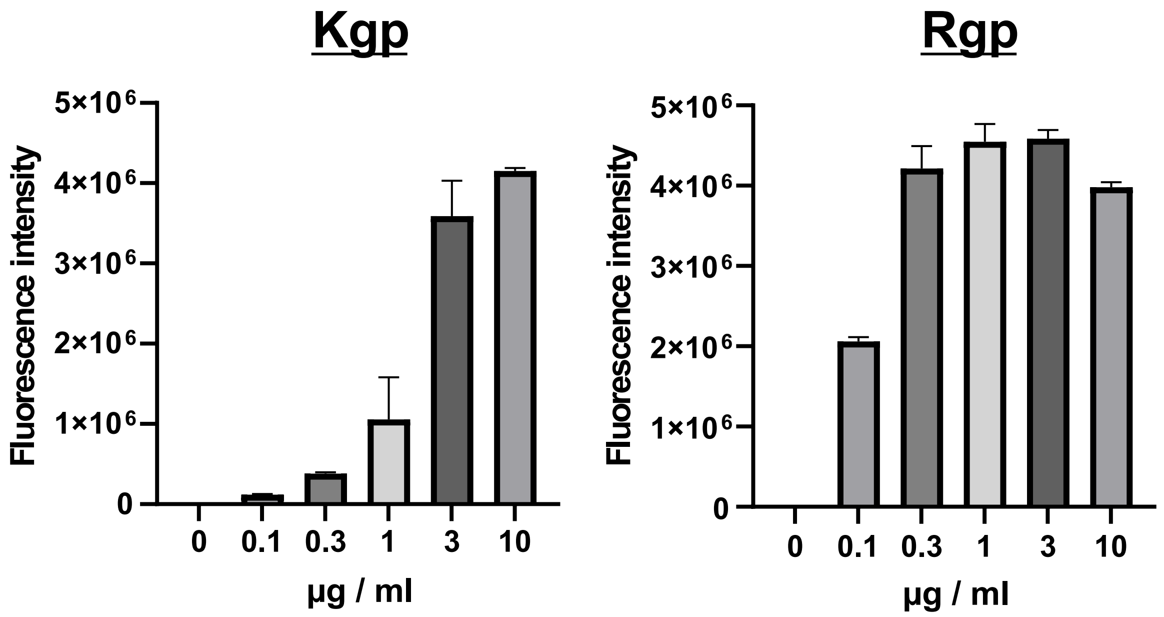


**Figure S1. Concentration-dependent gingipain activity in PCS.**
Lysine-specific gingipain (Kgp) and arginine-specific gingipain (Rgp) activities in PCS were measured using fluorogenic substrate assays. PCS was tested at the indicated concentrations, and enzymatic activity is shown as fluorescence intensity. Data are presented as the mean ± SEM; n = 3 independent measurements.

### Figure.S2

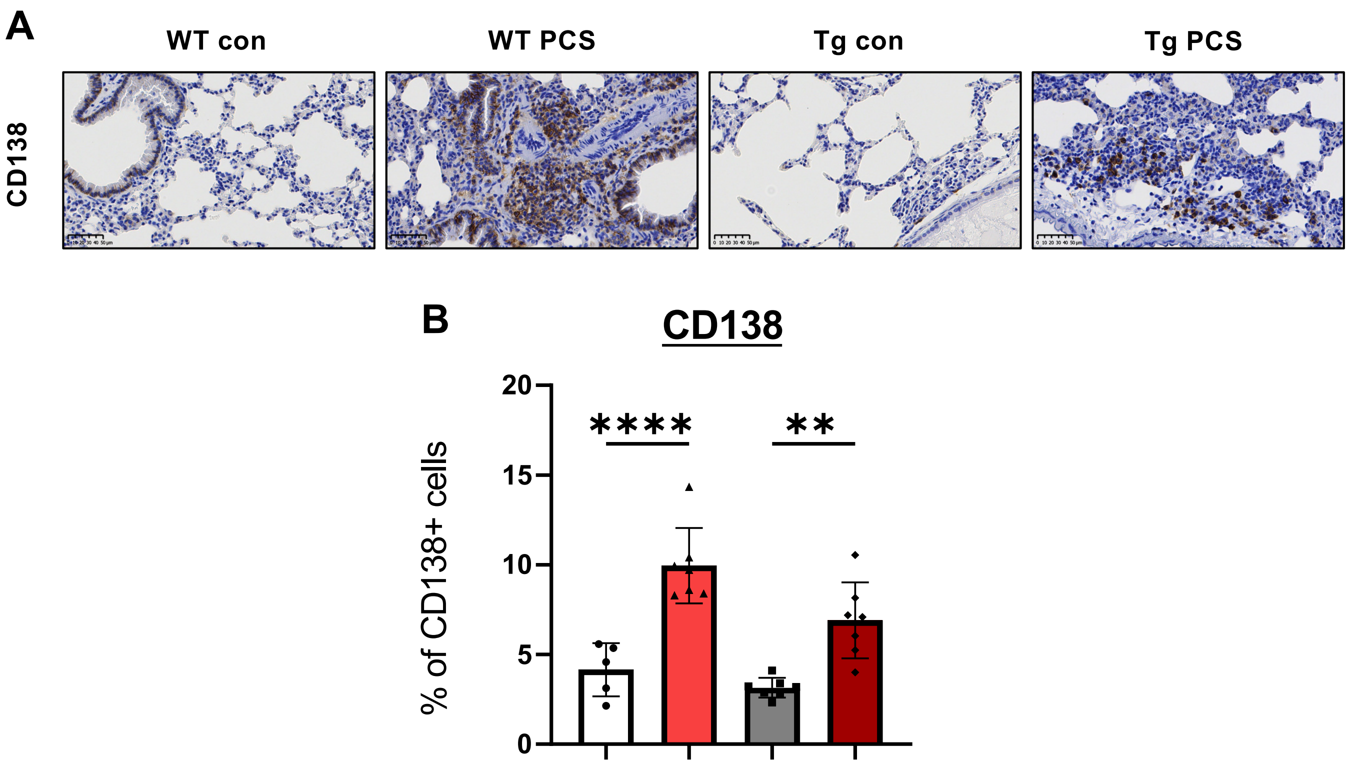


**Figure S2. Intratracheal administration of PCS induces airway-centered CD138-positive cell accumulation in WT and βENaC-Tg mice.**

(A) Representative immunohistochemical staining for CD138 in lung sections from each group. (B) Quantification of CD138-positive cells expressed as the percentage of marker-positive cells among total cells in the analyzed lung area. Quantification was performed using HALO image analysis software. Data are presented as the mean ± SEM; n = 5–7 mice per group. P values were assessed by one-way ANOVA followed by the Tukey-Kramer multiple-comparison test. **p < 0.01, ****p < 0.0001.

#### Figure.S3

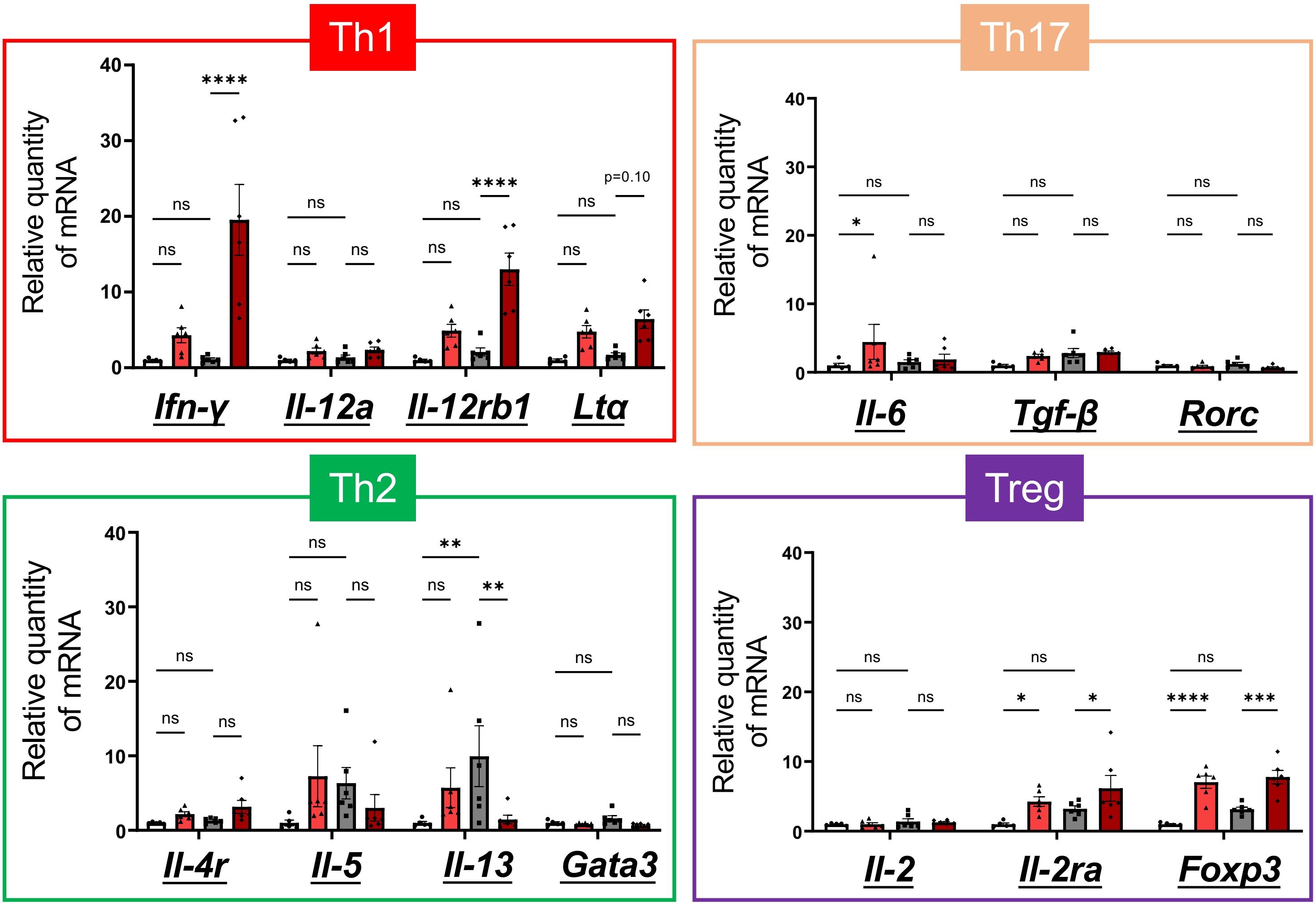


**Figure S3. PCS preferentially enhances Th1-associated gene expression in βENaC-Tg mouse lungs.**
Relative mRNA expression of T-cell differentiation-related cytokines and transcription factors was measured by RT-qPCR in lung tissues from WT control, WT PCS-treated, βENaC-Tg control, and βENaC-Tg PCS-treated mice. Genes associated with Th1 responses (*Ifng*, *Il12a*, *Il12rb1*, and *Lta*), Th17 responses (*Il6*, *Tgfb*, and *Rorc*), Th2 responses (*Il4r*, *Il5*, Il13, and *Gata3*), and regulatory T-cell responses (*Il2*, *Il2ra*, and *Foxp3*) were analyzed. Data are presented as the mean ± SEM; n = 5–7 mice per group. P values were assessed by one-way ANOVA followed by the Tukey-Kramer multiple-comparison test. *p < 0.05, **p < 0.01, ***p < 0.001, and ****p < 0.0001; ns, not significant. Exact p values are shown where applicable.

### Figure.S4

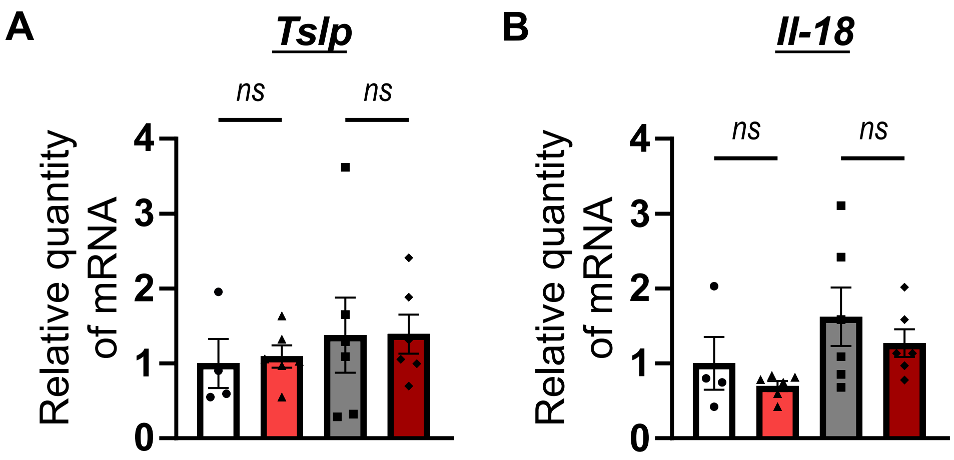


**Figure S4. PCS does not significantly induce epithelial cytokines associated with γδ T-cell activation in βENaC-Tg mouse lungs.** Relative mRNA expression of *Tslp* (A) and *Il18* (B) was measured by RT-qPCR in lung tissues from WT control, WT PCS-treated, βENaC-Tg control, and βENaC-Tg PCS-treated mice. Data are presented as the mean ± SEM; n = 5–7 mice per group. P values were assessed by one-way ANOVA followed by the Tukey-Kramer multiple-comparison test. ns, not significant.
